# Cerebrovascular Single-Nucleus RNA-Seq Reveals Heat Shock Activation and Vascular Remodeling in Alzheimer’s Disease and Primary Tauopathies

**DOI:** 10.64898/2026.06.26.734052

**Authors:** LD Estrella, S Dasgupta, A Gundavelli, H Li, HS Yang, S Chancellor, T Pastika, A Abdourahman, J Tamm, K Yanamandra, N Romanul, F Liao, K Zhao, G Lin, PP Srinivasa, X Wang, A Martin, E Asque, A Doering, JS Ried, RV Talanian, T Kwon, ME Woodbury, YY Grinberg, E Agastra, DH Oakley, BT Hyman, A Serrano-Pozo, T Zwang, S Das, RE Bennett

## Abstract

Cerebrovascular alterations are widely observed in both Alzheimer’s Disease (AD) and primary tauopathies. Here, we hypothesized that mechanisms of cerebrovascular alterations are shared between AD and primary tauopathies. We performed single-nucleus RNA sequencing of *postmortem* human inferior temporal gyrus to characterize transcriptomic changes across cerebrovascular cell types in AD and primary tauopathies (Corticobasal Degeneration, Pick’s disease, and Progressive Supranuclear Palsy). Differential gene expression analyses revealed disease-specific transcriptional programs across vascular cell populations. However, genes involved in the heat-shock response were consistently upregulated across all diseases, suggesting a conserved cerebrovascular stress response during neurodegeneration. We further identified marked cerebrovascular remodeling in AD relative to primary tauopathies, along with dysregulation of genes mapping to AD risk loci in endothelial cells. Transcriptomic findings were validated using tissue clearing, light-sheet microscopy, and immunofluorescence quantification of vascular markers. These results define a conserved vascular stress program alongside AD-specific remodeling, highlighting the vasculature as a therapeutic target in neurodegeneration.

## Introduction

Alzheimer’s disease (AD) and primary tauopathies are both characterized by the progressive accumulation of protein aggregates, the loss of neurons, and cognitive decline. Central to these disorders is the microtubule-associated protein tau, which undergoes post-translational modifications that promote oligomerization and the formation of intracellular inclusions. These inclusions acquire post-translational modifications and oligomerize to form intracellular deposits in each of these diseases^1,2^. Despite this shared tau pathology, the broader protein aggregation landscape differs across diseases. In AD, amyloid β plaques form in the extracellular space and tau forms distinct fibrillar tangles within neurons, particularly in brain regions related to memory and learning. In contrast, primary tauopathies are defined by tau pathology in the relative absence of amyloid β plaque deposition and exhibit substantial heterogeneity in their cellular and isoform-specific patterns. Primary tauopathies include Corticobasal Degeneration (CBD), Pick’s disease (PiD), and Progressive Supranuclear Palsy (PSP). In CBD and PSP, tau inclusions made up of mostly 4R tau isoforms accumulate in neurons, astrocytes, and oligodendrocytes. By comparison, Pick’s disease is characterized by round 3R tau inclusions in neurons and oligodendrocytes and profound cortical atrophy.

While the accumulation of protein aggregates in neurons and glial cells neuropathologically define these diseases, growing evidence indicates that other cell types in the brain play a critical role in the disease processes – notably in the cerebrovasculature. For instance, in AD, neuroimaging has revealed that reduced vascular perfusion to affected brain areas may be one of the earliest indicators of disease, with other studies indicating that blood-brain barrier (BBB) permeability is also impacted^3–6^. These findings, however, are not specific to AD, as blood flow abnormalities have also been reported in other dementias^7,8^. Interestingly, reductions in perfusion are strongly associated with the accumulation of tau pathology, suggesting a link between pathological protein deposition and altered brain vascular functions^9–11^.

To better understand this link, prior work from our group characterized the changes taking place in endothelial cells along the stereotypical spatiotemporal progression of AD using single nucleus RNA sequencing (snRNA-seq)^12^. We found a significant and unexpected upregulation of heat shock protein family genes, among other changes that were associated with disease stage. Building on these findings, we sought to determine whether this gene expression signature is specific to AD or represents a broader cerebrovascular response across tauopathies. We hypothesized that a substantial fraction of vascular transcriptional changes would be shared between AD and primary tauopathies, with a smaller subset being disease-specific. To test this, we compared AD-associated changes to those observed in primary tauopathies (CBD, PiD, PSP), thereby enabling us to disentangle the contribution of amyloid beta pathology to vascular alterations observed in AD. We expanded our analysis to include all vascular-associated cell types, including smooth muscle cells, pericytes, perivascular macrophages (PVMs), peripheral immune cells, and fibroblasts. Understanding these shared and disease-specific changes has the potential to provide novel insights into disease progression and identify new targets for diagnosis or treatment of AD and primary tauopathies.

## Methods

### Donor Tissues

All donor tissues were obtained from the Massachusetts Alzheimer’s Disease Research Center (MADRC) and were collected with approval from the Institutional Review Board (MassGeneral Brigham protocol 1999P009556) and consent of the next of kin. Disease status was based on detailed neuropathological analysis conducted by the MADRC Neuropathology Core. Frozen inferior temporal cortex (ITG, Brodmann Area 20) was used for all snRNA-seq experiments. Formalin-fixed paraffin embedded tissues from the contralateral hemisphere were used for histology and cleared tissue experiments, except 3D6 immunohistochemistry, which was done from cryostat sections. The brain area that was selected had minimal cerebral amyloid angiopathy (CAA), based on *postmortem* neuropathological reports and previously established scoring system^13^. The cohort included three donors with CAA score = 2, two donors with CAA score = 3, and one donor with CAA score = 4. All other donors had CAA scores = 0/NA. Donor samples were derived from two cohorts: an AD cohort^12,14^ and a tauopathy cohort (CBD, PiD, and PSP). Five control donors were shared across cohorts, with two additional controls unique to the AD cohort. In total, this study included *n* = 7 healthy aged controls (CTRL), *n* = 16 AD, *n* = 7 CBD, *n* = 8 Pick’s disease, and *n* = 33 PSP samples.

### Single-nucleus (sn)RNA-seq Analyses

#### Sample Preparation

For snRNA-seq, frozen cryostat sections were first prepared from ITG tissue blocks (**Figure 1A**). All tissues were assessed for RNA quality and had average RIN values > 3.8 (Table 1). Next, adjacent sections were subjected to nuclei extraction, labeling with antibodies to NeuN and Olig2 and were sorted to obtain a fraction depleted of neurons and oligodendrocytes, as previously described^12^. The sorted nuclei were then used to prepare cDNA libraries using the Chromium Single-Cell 3’ Reagents Kit V3 according to the manufacturer’s protocol (10x Genomics). Sequencing was then performed using an Illumina HiSeq 2000 on pooled samples to a depth of 30K reads per cell.

**Figure 1.**
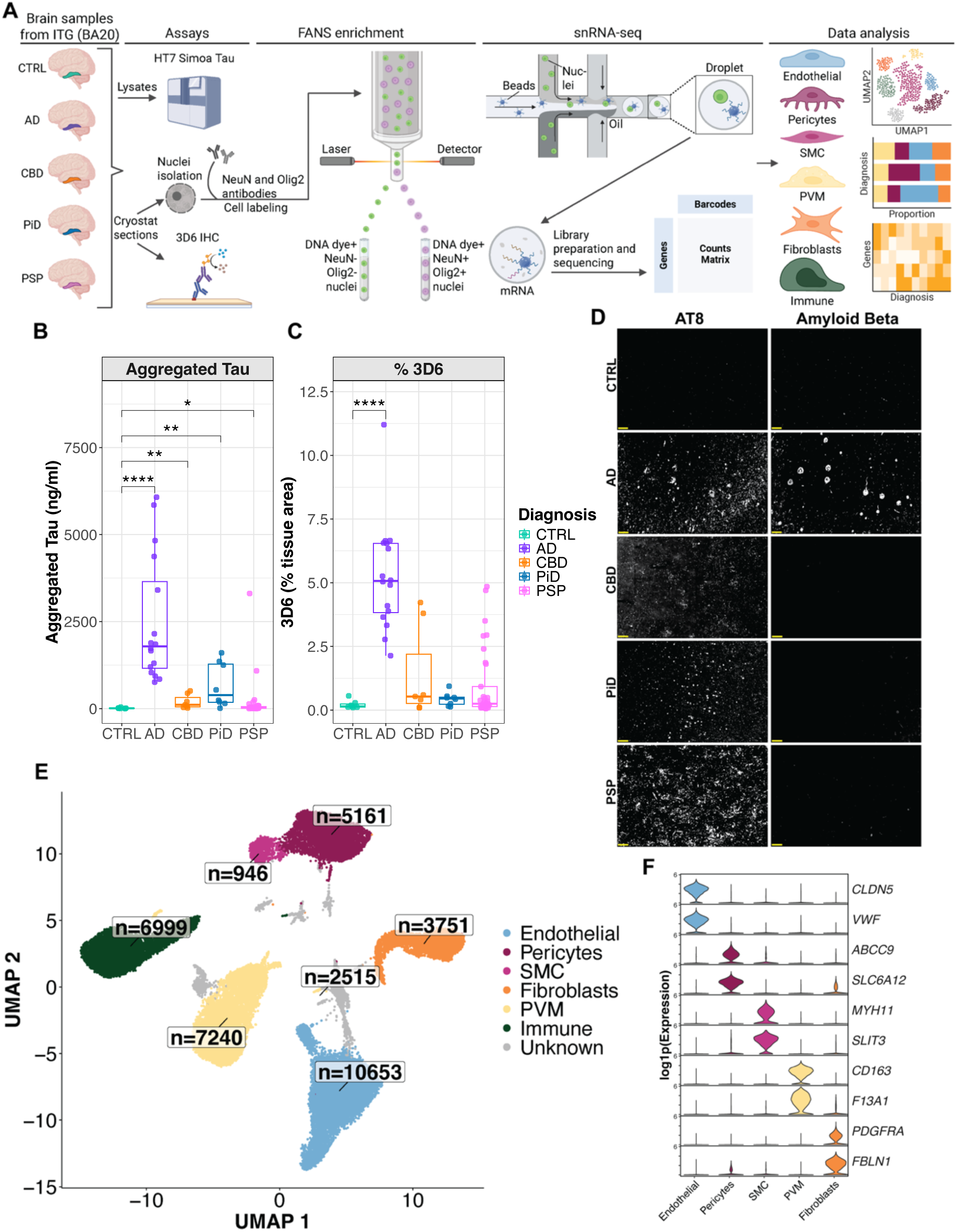
Single-cell SnRNA-seq workflow and characterization of human tissues. A) Illustration of sample processing and project workflow used for experimental assays and single-cell RNA-seq on human ITG (BA20) samples. Created in Biorender.com. B) Graphical quantifications from HT7 Simoa Tau assay using dissected ITG (BA20) human brain tissue lysates. C) Quantification of amyloid beta 3D6 immunohistochemistry from ITG (BA20) cryostat human brain tissue sections. Comparisons made using Kruskal-Wallis rank-sum test with Dunn’s post hoc analysis; \**p* ≤ 05, \*\**p* ≤ 01, \*\*\*\**p* ≤ 0001. D) Representative immunofluorescent images from contralateral ITG (BA20) tissue stained with AT8 (pTauSer202/Thr205, left) and amyloid beta (right) for neuropathological validation. Scale bars = 100 µm. E) UMAP visualization of the identified vascular-associated cell clusters, each containing their corresponding number of identified nuclei. F) Violin plots illustrating the expression levels of cell type-specific marker genes in NEUN^−^/OLIG2^−^ nuclei used to identify vascular-associated cells.

**Table 1.** Donor Summary Demographics. . Abbreviations: PMI – Postmortem interval, RIN – RNA integrity number, MAPT – Microtubule-associated protein tau haplotype, ADNC – Alzheimer’s disease neuropathological change score, s.d. – Standard deviation, NA – Not applicable or no available pathological examination.

| Summary | CTRL (*n* = 7) | AD (*n* = 16) | CBD (*n* = 7) | PiD (*n* = 8) | PSP (*n* = 33) |
| --- | --- | --- | --- | --- | --- |
| Age at Death<br>(mean ± s.d.) | 80 ± 12 | 81 ± 9 | 66 ± 6 | 65 ± 16 | 72 ± 7 |
| Sex<br>n (%female) | 4 (57.1) | 10 (62.5) | 2 (28.5) | 3 (37.5) | 16 (48.5) |
| PMI<br>(mean ± s.d.) | 16.8 ± 6.8 | 17.0 ± 9.5 | 24.6 ± 13.1 | 20.4 ± 10.9 | 18.9 ± 14.3 |
| APOE ε4<br>n (%) | 0 (0) | 7 (43.8) | 1 (14.3) | 2 (25.0) | 6 (18.2) |
| CERAD<br>NP Scores | None | Moderate<br>or<br>Frequent | None | None | None |
| MAPT<br>n (% H2) | 1 (14.3) | NA | 3 (42.9) | 2 (25.0) | 3 (9.1) |
| ADNC_A<br>n (Score) | 7 (0) | 3 (2)<br>13 (3) | 3 (NA)<br>4 (0) | 7 (NA)<br>1 (0) | 23 (NA)<br>8 (0)<br>1 (1)<br>1 (2) |
| ADNC_B<br>n (Score) | 7 (1) | 16 (3) | 3 (NA)<br>3 (0)<br>1 (2) | 7 (NA)<br>1 (0) | 17 (NA)<br>10 (0)<br>5 (1)<br>1 (2) |
| ADNC_C<br>n (Score) | 6 (0)<br>1 (1) | 8 (2)<br>8 (3) | 2 (NA)<br>5 (0) | 5 (0)<br>3 (NA) | 18 (NA)<br>15 (0) |
| RIN<br>(mean ± s.d.) | 6.0 ± 0.5 | 4.7 ± 1.6 | 4.4 ± 0.9 | 3.9 ± 0.9 | 4.8 ± 1.0 |

#### Data processing and quality control

Initial processing of raw reads was conducted using Cell Ranger (10x Genomics)^15^ with default settings, pre-mRNA package and aligned to the human GRCh38 genome. Seurat^16^ R package v5.0.0 was used for data processing, integration and clustering. Cells were subject to quality control and filtering to have greater than 800 detected genes (nFeature_RNA ≥ 800) but less than 6,000 detected genes (nFeature_RNA ≤ 6,000), less than 25,000 UMIs (nCount_RNA < 25,000) and less than 10% mitochondrial gene expression (percent.mt < 10). Subsequently, all mitochondrial genes were removed to avoid confounding downstream analyses. Doublets were identified and removed using the Python-based package Scrublet^17^ v0.2.3 with the doublet score threshold set to the 98th percentile. The filtered gene expression data were log1p-normalized using NormalizeData() and the top 2000 highly variable genes were identified for each donor using FindVariableFeatures() with selection.method = “vst”. The data were then scaled and centered across cells for each gene using ScaleData(). Dimensionality reduction was performed with RunPCA() and sample effects were accounted for via integration. Integration was carried out using RunHarmony(), leveraging the top 30 principal components and correcting for sample identity. This integration step was used to correct for batch effects arising from the inclusion of samples across the two cohorts. Uniform manifold approximation and projection (UMAP) for non-linear dimensionality reduction was done using RunUMAP() with the first 15 dimensions. The data were visualized in 2D space using DimPlot() with reduction = “umap”. Graph-based nearest neighbor clustering was performed using FindNeighbors() with reduction = “harmony” and dims = 1:15, followed by FindClusters() at resolutions 0.1 through 0.5. Resolution 0.1 was selected based on canonical marker expression (feature and violin plots), as it yielded clear separation of biologically distinct cell types without over-clustering. Cell types were subsequently annotated using these markers, identifying endothelial cells, pericytes, smooth muscle cells, PVMs, fibroblasts, and immune cells. Endothelial cells were subset from the Seurat object containing all cell types and processed using the same workflow. Resolution 0.1 was selected to classify them into capillary, venous, and arterial endothelial cells.

#### Cell type proportions

The proportion of each cell type was calculated within each diagnosis as the number of cells of that type divided by the total number of cells in that diagnosis. Donor-level cell-type proportions for each diagnosis were calculated as the number of cells of that type in a donor divided by the total number of cells in that donor. Differences in proportion across the diagnoses were assessed using Kruskal-Wallis tests, followed by Dunn’s post hoc tests with Benjamini–Hochberg (BH) correction for multiple comparisons. Bar plots were generated using the ggplot2 v4.0.1 package.

#### Differential expression analysis

For each cell type, disease group versus control (AD *vs.* CTRL, CBD *vs.* CTRL, PiD *vs.* CTRL, and PSP *vs.* CTRL) differential expression analysis was performed using FindMarkers() and the model-based analysis of single-cell transcriptomics (MAST) algorithm as done in previous AD single-cell sequencing studies^18–20^. Donor ID was included as a latent variable in the model. Differentially expressed genes (DEGs) were retained if the gene was detected in at least 5% of nuclei, if the absolute average log2FC > 0.25, and if the BH adjusted *P* value < 0.05. Overlap of DEGs across disease group versus control comparison was summarized using the ComplexUpset^21^ v1.3.3 package for each cell type.

#### Functional enrichment analysis

Functional enrichment of the significantly upregulated and downregulated DEGs was examined using the Gene Ontology (GO) database via enrichGO() from ClusterProfiler^22^ v4.16.0. Semantically redundant GO terms were collapsed using simplify()^23^ with similarity cutoff = 0.7. Six pathway categories of interest were defined from Gene Ontology (GO) enrichment results. For each disease *vs.* control comparison, collapsed GO outputs were filtered using pathway-specific keywords. Enriched terms were ranked by *P* value, and a representative pathway was selected for each pathway category and disease comparison based on statistical significance and biological relevance. Gene sets for each category were defined as the union of genes corresponding to the selected pathways across all disease comparisons, followed by manual curation to retain the most biologically relevant genes. Pathway-level results were visualized using bar plots.

#### Protein-to-gene expression correlations

Donor-level gene expression was computed using Seurat’s AverageExpression() on log-normalized counts. Spearman rank correlation was used to assess the association between gene expression and corresponding protein levels for selected genes. Correlation coefficients and associated *P* values were calculated using the cor.test() function in R. Results were visualized using scatter plots with fitted linear trend lines.

#### AD GWAS enrichment

AD GWAS genes reported by Yang *et al.* present in our single-nucleus RNA-seq dataset were retained for downstream analyses. To assess cell type–specific enrichment for these genes, average gene expression was computed across cells within each cell type using Seurat’s AverageExpression() on log-normalized counts. The resulting mean expression values were log1p-transformed, and gene-wise z-scores were calculated across cell types. The top 40 genes with the highest z-scores in capillary endothelial cells were selected and visualized using the ComplexHeatmap v2.24.1 package. The overlap between AD GWAS genes and DEGs in capillary endothelial cells (determined by MAST and filtered using previously mentioned criteria) was determined for each disease *vs.* control comparison. Genes significant by BH adjusted *P* value in at least one disease *vs.* control comparison were retained, and the number of overlapping genes per disease was summarized using bar plots. Subsequently, these genes were ordered based on the mean scaled log2 fold change across all diagnoses, where log2 fold change values were derived from the MAST differential expression analysis. To enable symmetric visualization of up- and downregulation, log2 fold changes were transformed as 2^avg_log2FC for positive values and −(1/2^avg_log2FC) for negative values.

### Donor Genotyping

Genomic DNA was extracted from cerebellum samples corresponding to the same donors used for snRNA-seq. Cerebellum samples were disrupted and homogenized by beadmill with the TissueLyzer 2 (Qiagen, 85300) following the homogenization strategy at a frequency of 30 Hz for 30 sec. Cerebellum lysates then passed through QIAshredder columns (Qiagen, 79656) and were centrifuged per manufacturer’s instructions at top speed (20,000 xg) for 2 minutes. Homogenized lysates then followed the DNA extraction workflow on the Qiagen QIAcube HT (Qiagen, 9001896) with the QIAamp DNA mini kit chemistry (Qiagen, 51306) following the manufacturer’s instructions. DNA eluate was assayed for quality with fluorescent quantification using the Qubit 1x dsDNA kit (Thermofisher Scientific, Q32853) and automated gel electrophoresis using the TapeStation with the genomic DNA screentape assay (Agilent, 5067-5365) was performed following manufacturer’s instructions respectively. Genotype microarray data was then generated from 200 ng of extracted DNA. The DNA was processed through the Illumina Infinium LCG workflow following manufacturer’s instructions and genotype data collected from the Global Diversity Array v1 with the iScan and Autoloader (Illumina, 20031669, SY-101-1001, PCE4-EX respectively). Call rate quality control (passing at or over 0.99) was assessed with Genome Studio v2.0 (Illumina) before entering downstream analysis workflows.

### eQTL analysis

Donor-level gene expression was computed using Seurat’s AggregateExpression() on raw counts, followed by normalization and log1p transformation. For each gene, corresponding SNP identifiers were obtained, and genotype information was extracted from orthogonally collected genomic data to calculate allele dosage. Associations between gene expression and genotype dosage were evaluated using a linear model: Expression ∼ Dosage + Age + Sex + Diagnosis. Genes with nominally significant associations were visualized using boxplots of donor-level expression.

### Tau HT7-HT7 SIMOA

Single Molecule Array (SIMOA; Quanterix) bead-based tau aggregates assay was developed using a mouse anti-HT7 antibody (Thermo Fisher Scientific, RRID: AB_2314654) as both capture and detection. The assay was prepared according to the manufacturer’s protocol. Recombinant full length P301L tau aggregates were made as described^24^ and were used as a calibrator and included in each run to generate standard curve. HD-X instrument, buffers, helper beads and streptavidin B-galactosidase, and enzyme substrate resorufin Beta-D-galactopyranoside were obtained from Quanterix. Assays were performed according to manufacturer’s instructions. All samples were diluted in the Tau Calibrator Diluent (Quanterix).

### 3D6 DAB Immunohistochemistry

Cryosections (10 µm thickness) were taken from the same pieces used for snRNA-seq. For Amyloid-beta immunohistochemistry, PFA-post fixed frozen cryostat sections adjacent to those used for snRNA-seq were subjected to immunohistochemistry with mouse monoclonal anti N-terminal Amyloid-beta antibody clone 3D6 (2 µg/mL).

### Fluorescent Immunohistochemistry

FFPE tissue blocks containing ITG (BA20, contralateral to the transcriptomic samples) were cut coronally at the level of the lateral geniculate nucleus and were used for this analysis. Tissues were cut at 8 µm thickness on a microtome and mounted on Superfrost Plus slides (Fisher Scientific). Sections were rehydrated in xylene and descending concentrations of ethanol, followed by epitope retrieval using citrate buffer (pH 6.0, containing 0.05% Tween-20) and heat (20% microwave power for 20 minutes). For immunolabeling, sections were incubated in blocking buffer (5% BSA and 0.25% Triton-X in TBS) for 1 h at RT. After blocking, sections were incubated overnight at 4 C with the following primary antibodies: GLUT1 (1:500, EMD Millipore, cat #07-1401), HSP90AA1 (1:250, Invitrogen, cat #PA3-013), HLA-A (1:400, Abcam, cat #ab52922), and PLCG2 (1:250, ThermoFisher, cat # PA5-86233). Sections were then thoroughly washed with TBS and incubated with Alexa dye-conjugated secondary antibodies in blocking buffer for 1 h at RT. After secondary antibody incubation, another round of washing steps was performed prior to Trueblack (Biotum) application for 30 sec to quench autofluorescence. Slides were mounted with Fluoromount G – DAPI solution, coverslipped, and imaged using an Olympus VS120 slide scanner under a 20x magnification objective.

### Single-Cell Quantification of Fluorescent Immunohistochemistry Images

Slide scans were analyzed using QuPath software^25^ (Version 0.6.0) with the workflow described in **Supplemental Figure 1**. Here, an ROI with a consistent area was drawn within the ITG of each case. To achieve individual cell measurements similarly to the transcriptomics data, all DAPI-positive cells within the ROI were identified using QuPath’s cell detection tool. Following cell detection, an object classifier was applied to detect endothelial cells using GLUT1 intensity thresholds. The classification resulted in detection of DAPI-positive/Glut1-negative and DAPI-positive/Glut1-positive cell populations. The fluorescent intensity for the markers of interest (HSP90AA1, PLCG2, and HLA-A) was measured for each individual vascular cell that was identified. The reported intensity values were normalized to the ROI background intensity as follows:

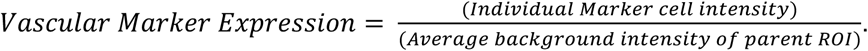

Analysis of all individual cell intensity values was performed using the repeated measures Linear Mixed Model (LMM) statistical test. The data was log-transformed to achieve normality. LMM test was used having diagnosis, age, and sex as fixed variables and donor ID as a random effect variable. After each statistical test, the performance was validated by assessing the model’s linearity, homogeneity of variance, and the normality of random effects.

### 3D Imaging of Blood Vessels

#### Tissue clearing

Formalin-fixed human ITG samples from a subset of donors included in the snRNA-seq study (contralateral side) were used for tissue clearing experiments to evaluate vasculature (**Supplemental Figure 2**). This included samples from n = 2 healthy controls, n = 4 AD, n = 2 PSP, n = 1 PiD, and n = 2 CBD donor cases, which underwent tissue clearing procedures similarly to those described previously^26,27^.

To remove all formalin, brain samples were thoroughly washed in PBS for at least 24 h at RT with constant solution changes. Tissue samples were then transferred to individual 35 mm Petri dishes and embedded in a gel block by pouring warm 3% agarose gel solution in PBS over the tissue. The gel was then cooled to solidify and cut into a block to provide rigidity for cutting even slices. The tissue was secured on a vibratome (Leica Biosystems, VT1000 S Vibrating Blade Microtome) using super glue and 1 mm-thick sections were cut for each case. The agarose was removed from each section through gentle manipulation with blunt forceps and paintbrushes prior to starting the tissue clearing protocol.

Brain sections were each incubated in 5 mL of SHIELD OFF solution composed of 50% SHIELD-Epoxy buffer (LifeCanvas Technologies), 25% SHIELD Buffer (LifeCanvas Technologies), and 25% distilled water for 24 h at 4 °C. Samples were then transferred to SHIELD ON solution (5 mL/tube) containing 7:1 SHIELD ON Buffer : SHIELD-Epoxy buffer (LifeCanvas Technologies) for 5 h at 37 °C wih light shaking, followed by overnight incubation with 100% SHIELD ON buffer under the same conditions. After crosslinking steps, sections were thoroughly washed four times for 10 min with PBS at RT while shaking. Sections were then incubated in 8-10 mL of delipidation buffer (LifeCanvas Technologies) for 24 h at 37 °C while shaking.

After delipidation, sections underwent epitope retrieval through incubation in 8-10 mL of 0.1M Glycine for 1 h at RT with light shaking. After this, sections were transferred to 8-10 mL of PBS and placed in an oven at 70 °C for 1 h. Following epitope retrieval steps, sections were washed three times for 10 min with PBS, kept in PBS, and exposed to high-intensity LED white light (∼250 cm separation) for 3-4 h to quench tissue autofluorescence. Tissue samples were then thoroughly washed and incubated for 5 days at RT with GLUT1 (1:500, EMD Millipore, cat #07-1401 conjugated to Alexa fluor 647) diluted in PBS-T (0.25% Tween 20).

After immunolabeling, samples were thoroughly washed with PBS and later incubated in 50% DI water and 50% Easy Index (LifeCanvas Technologies) overnight at RT. Following this, samples were incubated in 100% Easy Index for at least 4 h at RT. Cleared tissues were placed on SuperFrost microscope slides and embedded by pouring agarose (2% in Easy Index) directly on the tissues.

#### Light-sheet microscope imaging

Cleared tissues were imaged using LifeCanvas Technologies megaSPIM light sheet microscope equipped with a 3.6x lens objective under the 639 nm laser channel. Captures from the light sheet microscope were stitched at a voxel size of 1.826 x 1.826 x 1.826 µm. Stitched images were then converted into Imaris files for visualization and image quality assessment.

#### Generation of a Vessel Segmentation Mask

To facilitate computing processing, the generated Imaris file was downsized on the x and y planes by half, then converted into an OMEzarr file in order to be imported into Ilastik software and generate a vessel mask using Ilastik’s pixel classifier. Using all 37 available features, Ilastik’s pixel classifier was trained to detect positive and negative pixels and trace vessels across all z-stacks. Once the training was validated, all z-stacks were processed, and a simple segmentation mask was exported as an HDF5 file format. MATLAB (version R2022a 9.12.0) was then used to extract only the positive signal of the traced vessels and export this as a TIFF file. ImageJ was used to visualize the exported vessel segmentation mask and confirm accurate vessel tracing. The mask was converted into a binary image and the binary feature “Fill Holes” was used. The resulting z-series was exported as an OME-TIFF file, converted back into an Imaris file, and resized back to its original dimensions.

#### Imaris Filament Tracer and Vessel Quantification

Using Imaris’ Filament Tracer feature (version 10.2.0), two cubic ROIs (∼4 mm^3^ each) were placed on the brain gyrus covering predominantly gray matter. Parameters were set for multiscale filament tracing, setting the smallest diameter at 3 µm and largest diameter between 60-80 µm. Both filament tracing seed points and detected segments were trained, and a filter was set to remove segments with lengths < 10 µm. Filament Tracer output statistics were used to quantify vessel features like volume, number of segments, length, and straightness.

## Results

### Study Design and Neuropathological Characterization of Human Inferior Temporal Cortex Samples

To compare the gene expression signatures of vascular cells while avoiding region-specific differences, we selected a region of temporal cortex (ITG), as this brain area is commonly affected by tau aggregation in AD and primary tauopathies (**Figure 1A**)^28–31^. Using a tau single-antibody sandwich ELISA, the extent of oligomeric tau varied considerably with AD samples showing the highest values and healthy aged controls showing the lowest values, as expected (**Figure 1B**). While amyloid β accumulation in plaques is a defining feature of AD, some CBD and PSP individuals also had areas of amyloid β positivity in ITG (**Figure 1C**), but had CERAD scores of none or sparse, indicating the absence of significant neuritic plaque pathology. In addition, histopathological assessment of tau aggregation showed the expected pathological tau profiles representative of each disease, having neurofibrillary tangles in AD, astrocytic inclusions in CBD and PSP, and Pick bodies in PiD (**Figure 1D**). Only AD cases displayed profound amyloid β plaque accumulation in this cortical region (**Figure 1D**). A full description of sample donors included in this study is depicted in **Table 1**.

### Disease-Specific Cerebrovascular Remodeling in Alzheimer’s Disease and Primary Tauopathies

Across all samples, we were able to identify 37,265 vascular-associated cell nuclei including: endothelial cells, smooth muscle cells, pericytes, PVMs, fibroblasts, and peripheral immune cells (**Figure 1E**). Cell type was confirmed using previously published marker genes^18,32^ (**Figure 1F**). Endothelial cells were further categorized by zonation (arterial, capillary, or venous), with capillary endothelial cells comprising most of the population, as previously found using this analysis pipeline (**Supplemental Figure 3**). Surprisingly, capillary endothelial cells were underrepresented in all primary tauopathies compared to the proportion observed in AD. By comparison, there was a slight trend towards an overrepresentation of capillary endothelial cells in AD samples compared to normal aging controls (**Figure 2A & 2B**). Other cell types such as pericytes and PVMs did not follow this pattern. For instance, pericytes showed a significant overrepresentation in the primary tauopathies (CBD and PSP) compared to AD (**Figure 2A & 2C**), while PVMs showed significant overrepresentation in Pick’s disease and PSP, compared to AD (**Figure 2A & 2D**).

**Figure 2.**
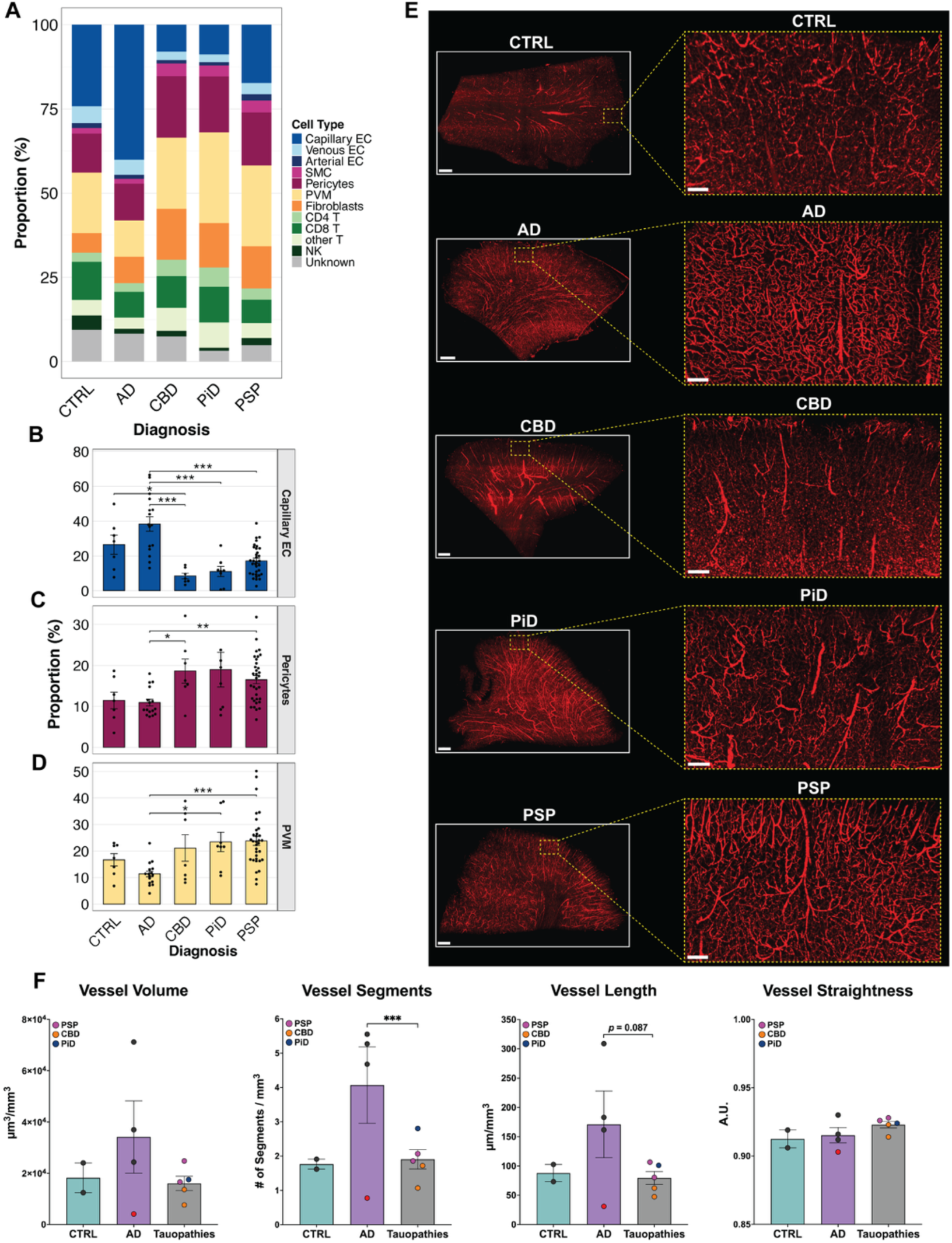
Cell proportion changes and disease-specific vascular remodeling in AD and primary tauopathies. A) Stacked bar chart demonstrating the percent contribution of the identified cell types to the total number of identified nuclei within each diagnosis. B) – D) Individual bar plots displaying the percent number of capillary endothelial cells (Capillary ECs, B), pericytes (C), and perivascular macrophages (PVMs, D) in each donor to the total number of identified nuclei within that donor. Bars represent the mean proportion across donors within each diagnosis group, and points indicate individual donors. Error bars denote SEM. Overall differences across diagnosis groups were assessed using a Kruskal–Wallis rank-sum test. When significant, post hoc pairwise Dunn’s tests were performed, with Benjamini–Hochberg correction for multiple comparisons. Significance is denoted as adjusted *P* value < 0.05 (*), < 0.01 (**), and < 0.001 (***). E) Representative immunofluorescent images of 3D-reconstructed vascular networks from 1-mm thick tissue samples from contralateral ITG (BA20) stained with endothelial cell maker GLUT1 (red). Individual zoomed-in ROIs focus at cortical layers II/III grey matter. Whole tissue image scale bars = 1mm; zoomed-in ROI scale bars = 200 µm. F) Graphical quantifications of 3D vessel reconstruction characteristics including volume, length, segments, and straightness from control (n = 2), AD (n = 4), and primary tauopathy samples (total n = 5 including CBD n = 2, PiD n = 1, and PSP n = 2). One AD outlier sample was removed due to poor quality staining. Statistical comparisons between AD and primary tauopathies using two-tailed Welch’s t test. Statistical significance: *** *p* ≤ 001.

To verify and better understand the observed cerebrovascular proportion differences between AD and primary tauopathies, we utilized light sheet microscopy on cleared ITG tissues from the same donors used in the transcriptomic analyses (CTRL = 2, AD = 4, CBD = 2, PiD = 1, PSP = 2). A 3D vessel reconstruction and vessel mask was generated to visualize the vascular architecture and measure vessel features such as volume, segments, length, and straightness. Since the snRNA-seq proportion analyses showed that endothelial cell proportions are underrepresented in primary tauopathies, samples from CBD, PiD, and PSP donors were pooled and categorized as primary tauopathies. The cerebrovasculature of AD tissues qualitatively appeared to have increased vessel density and a more complex vessel branch distribution compared to control and primary tauopathies, where vascular networks were less elaborate – particularly in deeper cortical layers and white matter (**Figure 2E**). The quantification of vessel features showed that AD donors had a significant increase in the number of vessel segments and vessel length per volume, compared to controls and primary tauopathies (**Figure 2F**). Together, these data suggest that vascular remodeling differentially occurs in AD *vs.* primary tauopathies.

### Shared and Distinct Differential Gene Expression Changes Between Alzheimer’s Disease and Primary Tauopathies

We next performed a differential gene expression (DGE) analysis to compare changes taking place in each disease versus control. Significantly up- and down-regulated genes were observed in all vascular cell types in each disease (**Supplemental Figure 4**). However, clear differences were observed in the extent that each cell type appears to be impacted by disease. For example, capillary endothelial cells expressed the largest proportion of differentially expressed genes (DEGs) in AD and PSP (**Figure 3A; Supplemental Figure 4A & 4D**), while pericytes appear to be more affected in PiD and CBD (**Figure 3B; Supplemental Figure 4B & 4C**). Other cell types, such as fibroblasts in AD, CBD and PiD, PVMs in AD and CBD, and immune cells in CBD also showed significant gene expression differences (**Figure 3 C-D; Supplemental Figure 4 A-D)**. No significant DEGs were identified in smooth muscle cells. PSP was unique in that few DEGs were identified in vascular cell-types other than capillary endothelial cells.

**Figure 3.**
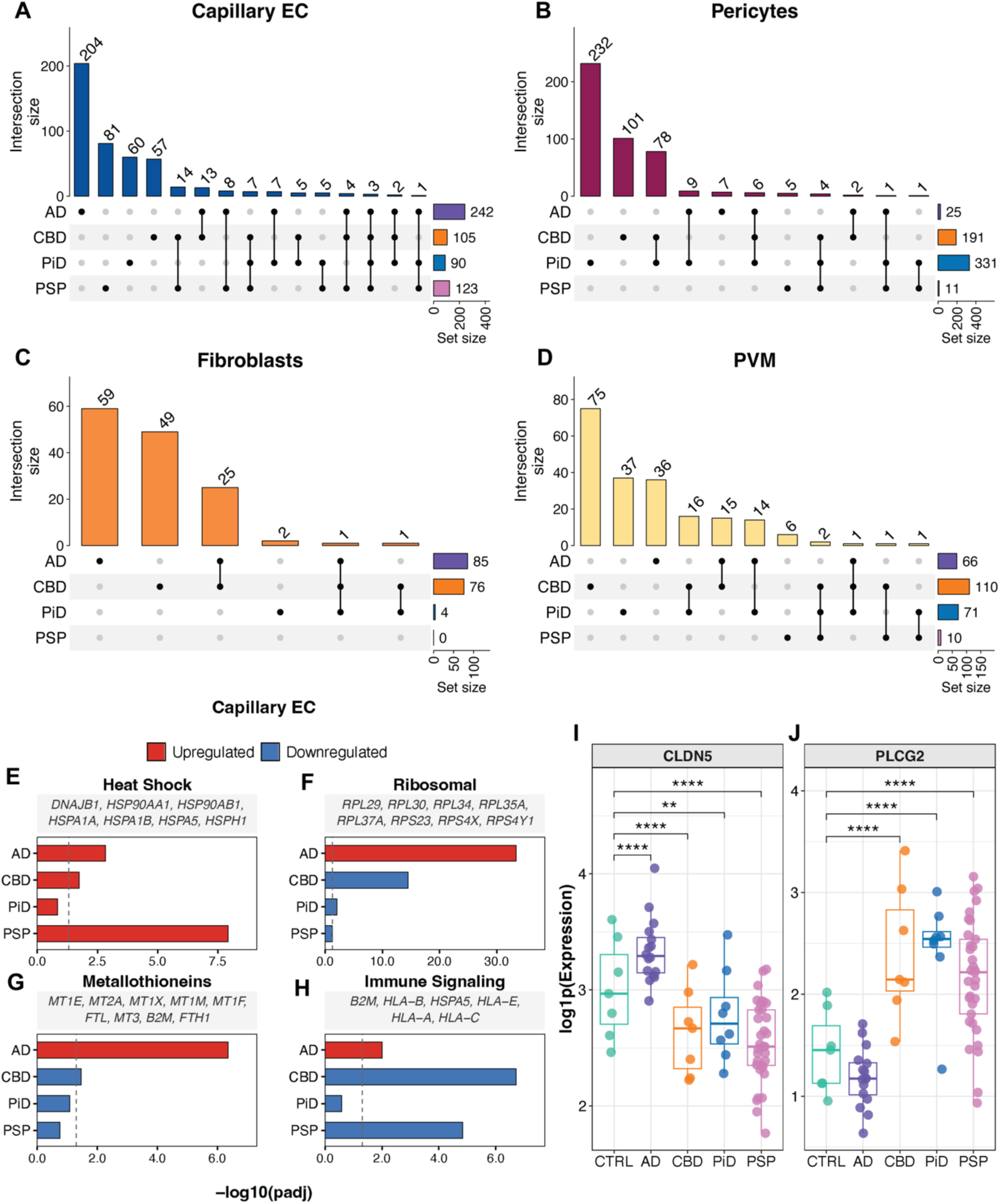
Analysis of differentially expressed genes (DEGs) in AD and primary tauopathies. A) – D) Upset plots displaying the number of DEGs for each pathological diagnosis vs control and whether DEGs are shared between such comparisons in capillary endothelial cells (Capillary ECs, A), pericytes (B), Fibroblasts (C), and perivascular macrophages (PVMs, D). E) – H) Representative bar plots from functional pathway analysis in capillary endothelial cells, showing pathway-level changes for each diagnosis relative to control across heat shock response (E), ribosomal processes (F), metallothionein (G), and immune signaling (H). Bar length represents −log10(adjusted *P* value) for each diagnosis vs control comparison. Bar color indicates the direction of the pathway (red, upregulation; blue, downregulation). The dashed vertical line marks the significance threshold at adjusted *P* value = 0.05. Gene labels above each panel show representative genes contributing to the pathway. I) – J) Box plot representations of donor-averaged log1p expression for CLDN5 (I) and PLCG2 (J) in capillary endothelial cells across all diagnoses. Each point represents a donor. Significance annotations correspond to MAST-derived adjusted *P* values from capillary endothelial cell differential expression analyses for each diagnosis vs. control comparison and are denoted as adjusted *P* value < 0.05 (*), < 0.01 (**), < 0.001 (***), and < 0.0001 (****).

Since capillary endothelial cells were commonly affected across disease states, we focused on shared up- or down-regulated genes in these cells. Only 3 differentially expressed genes (DEGs) were shared across all comparisons (AD *vs.* CTRL, CBD *vs.* CTRL, PiD *vs.* CTRL, and PSP *vs.* CTRL), including *CLDN5, HSP90AA1 and RPL3* (**Figure 3A**). In the case of primary tauopathies, 7 DEGs were shared across their respective comparisons with controls, including *CLIC1, H3F3B, MALAT1* and *PLCG2* in addition to *CLDN5, HSP90AA1* and *RPL3* (**Figure 3A**). No common genes were shared across all disease states in any other cell-type (Figure 3 B-D). In AD, *HSPA1A, HSPA1B, HSPH1, HSP90AA1*, among other heat shock protein genes, were highly significantly enriched with average log2 fold change values ranging from 1.66 to 2.28 (**Supplemental Table 1)**. While AD displayed the greatest enrichment of these genes, pathway analysis identified terms including protein folding and HSF1-transactivation (heat shock proteins), which were commonly enriched across disease states (**Figure 3E**). This includes the gene *HSP90AA1,* which was universally upregulated compared to controls. This was not unique to capillary endothelial cells. In pericytes, heat shock protein genes like *HSP90AA1* were also upregulated in PiD and PSP with log2 fold change values ranging from 0.87 to 0.91 (**Supplemental Table 1**). Altogether, these data highlight that, in the setting of tau pathology, vascular cells commonly upregulate heat shock pathway genes.

Similarly, several genes encoding proteins involved in ribosome biogenesis were significantly upregulated in AD but not in tauopathies. This upregulation included genes encoding both the small 40S (*RPS11, RPS13, RPS27, RPS27A*, etc.) and large 60S (*RPL3, RPL9, RPL23A, RPL41*, etc.) ribosomal subunits. Many of these genes were oppositely regulated in tauopathies, where they appeared to be less abundant. For example, *RPL3* was significantly downregulated in CBD, PiD, and PSP *vs.* controls (**Figure 3F**). Metallothionein genes, which we had previously observed to be upregulated in AD capillary endothelial cells^12^, were only increased in AD (**Figure 3G**). Notably, major histocompatibility complex type I protein coding genes, including *HLA-A-, HLA-B, HLA-C*, and *HLA-E* were all significantly downregulated in CBD (log2FC -0.49 to -1.30) and in PSP (log2FC -0.45 to -0.84). No significant differences were observed in these genes in PiD, and in AD an increase in *HLA-B* expression was noted (log2FC = 0.48) (**Figure 3H**). Among other gene expression changes that were unexpectedly different between AD and primary tauopathies included, in capillary endothelial cells, the gene for tight junction protein claudin 5 (*CLDN5*) which was significantly downregulated across all tauopathies but significantly upregulated in AD (**Figure 3I & Supplemental Table 1**). Interestingly, *PLCG2*, a gene encoding for phospholipase C gamma 2 and a known AD risk gene^18^, was significantly upregulated in primary tauopathies but unchanged in AD *vs.* controls (**Figure 3J**).

Lastly, to confirm that incidental amyloid β pathology was not skewing these results, we separated the PSP cohort into “low”- and “high”- amyloid β and compared each to control. DEGs identified in each group were nearly identical, indicating the observed gene expression changes are likely independent of amyloid β pathology (**Supplemental Figure 5**). In sum, AD-specific increases in *CLDN5*, immune signaling, ribosomes, and metallothioneins are not shared by tauopathies, while other changes such as increased *PLCG2* are unique to primary tauopathy.

### Heat Shock-Related Gene Changes Directly Translate into Protein Abundance Changes within Brain Endothelial Cells

To evaluate how the observed gene changes reflect protein expression differences, we performed immunofluorescent histology using FFPE sections from the contralateral hemisphere of a subset of donors (15 individuals, n = 3 per diagnosis). In line with gene expression data, we observed a significant increase of HSP90AA1 protein levels within the endothelial cells of AD, PiD, and PSP cases, when compared to controls (**Figure 4A & 4B**). HSP90AA1 in CBD cases did not reach statistical significance, though still trended towards increased protein expression (**Figure 4B**). Independent of diagnosis, the expression of *HSP90AA1* gene correlated significantly with the abundance of HSP90AA1 protein in endothelial cells (**Figure 4C**).

**Figure 4.**
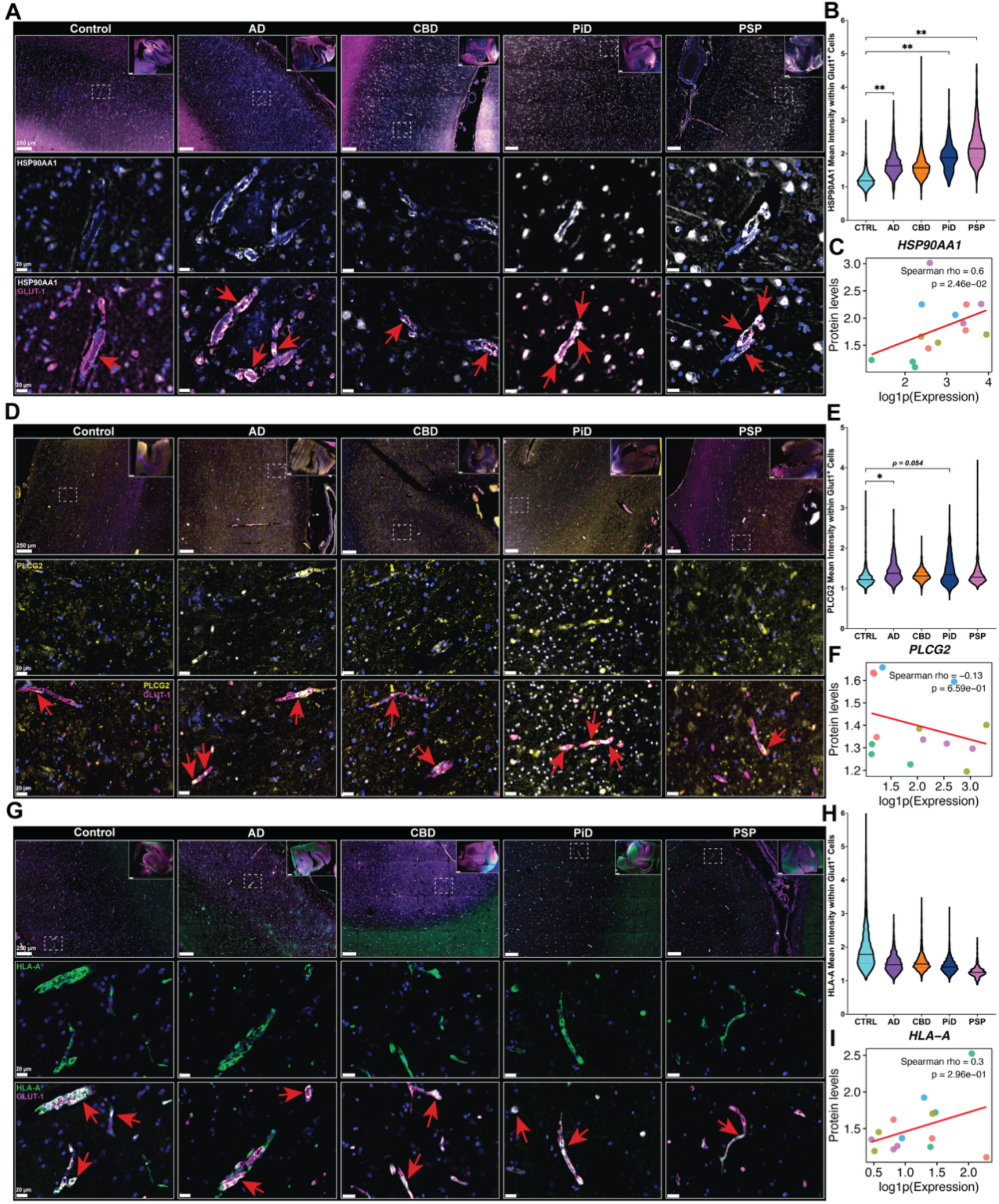
Cross-modal validation of vascular transcriptomic changes at the protein level. A) Representative immunofluorescent images of human ITG across diagnoses (BA20, contralateral to transcriptomic samples) stained for HSP90AA1 (white), endothelial cell marker GLUT1 (magenta), and nuclear counterstain DAPI. B) Quantification of HSP90AA1 mean fluorescent intensity within GLUT1^+^ endothelial cells across diagnoses. C, F, I) Spearman correlation analysis between donor-averaged log1p gene expression (snRNA-seq) and corresponding protein expression (IHC) for HSP90AA1 (C), PLCG2 (F), and HLA-A (I) in brain endothelial cells, independent of diagnosis. Correlation strength was assessed using Spearman rank correlation, and the reported statistics indicate Spearman’s rho and the corresponding *P* value. Red lines represent the fitted linear trend. D) Representative immunofluorescent images stained for PLCG2 (yellow), GLUT1 (magenta), and DAPI. E) Quantification of PLCG2 mean fluorescent intensity within GLUT1^+^ endothelial cells across diagnoses. G) Representative immunofluorescent images stained for HLA-A (green), GLUT1 (magenta), and DAPI. H) Quantification of HLA-A mean fluorescent intensity within GLUT1^+^ endothelial cells across diagnoses. Slide overview scale bar = 2 mm. Red arrows correspond to vascular expression of the protein marker of interest. Fluorescence intensity was quantified within GLUT1⁺ endothelial cells as a proxy of vascular protein expression and normalized to each individual parent ROI background signal. IHC statistical comparisons across diagnoses were performed using a linear mixed-effects model with diagnosis, age, and sex as fixed effects and donor ID as a random effect. n = 3 donors per diagnosis. Statistical significance: *p ≤ 0.05; **p ≤ 0.01.

Other gene-protein correlations were less robust. *PLCG2*, for example, showed a transcriptomic upregulation in primary tauopathies (**Supplemental Table 1**), but histological assessment showed a significant increase in endothelial cell PLCG2 protein abundance in AD compared to controls with no change in tauopathies (**Figure 4D**, **E**). No correlation was found between our snRNA-seq and histological data (**Figure 4F**). Similarly, we assessed the expression of HLA-A protein in endothelial cells and, while expression trends were similar to gene expression patterns, these differences did not reach statistical significance (**Figure 4G-I**). In sum, while strong gene-protein relationships were observed in HSP90AA1, gene expression changes were not always reflected by changes at the protein level, as highlighted by prior reports^33,34^.

### AD GWAS Genes are Enriched in the Cerebrovasculature and are Differentially Expressed Between Alzheimer’s Disease and Primary Tauopathies

Previous studies have noted that genes identified by AD Genome-Wide-Association-Studies (GWAS) are expressed at high levels in both myeloid and endothelial cells, suggesting a key role in AD pathogenesis^18,35,36^. We similarly evaluated a list of 651 AD-risk genes to determine their expression across cells. Of these, 588 were expressed in capillary endothelial cells. In many cases, the expression was pronounced in capillary endothelial cells compared to other vascular cell types and, surprisingly, also to microglia. The top 40 capillary endothelial cell enriched GWAS genes are summarized in **Figure 5A** along with their relative enrichment in the other vascular cell types. This confirms that capillary endothelial cells express a substantial number of AD-risk genes.

**Figure 5.**
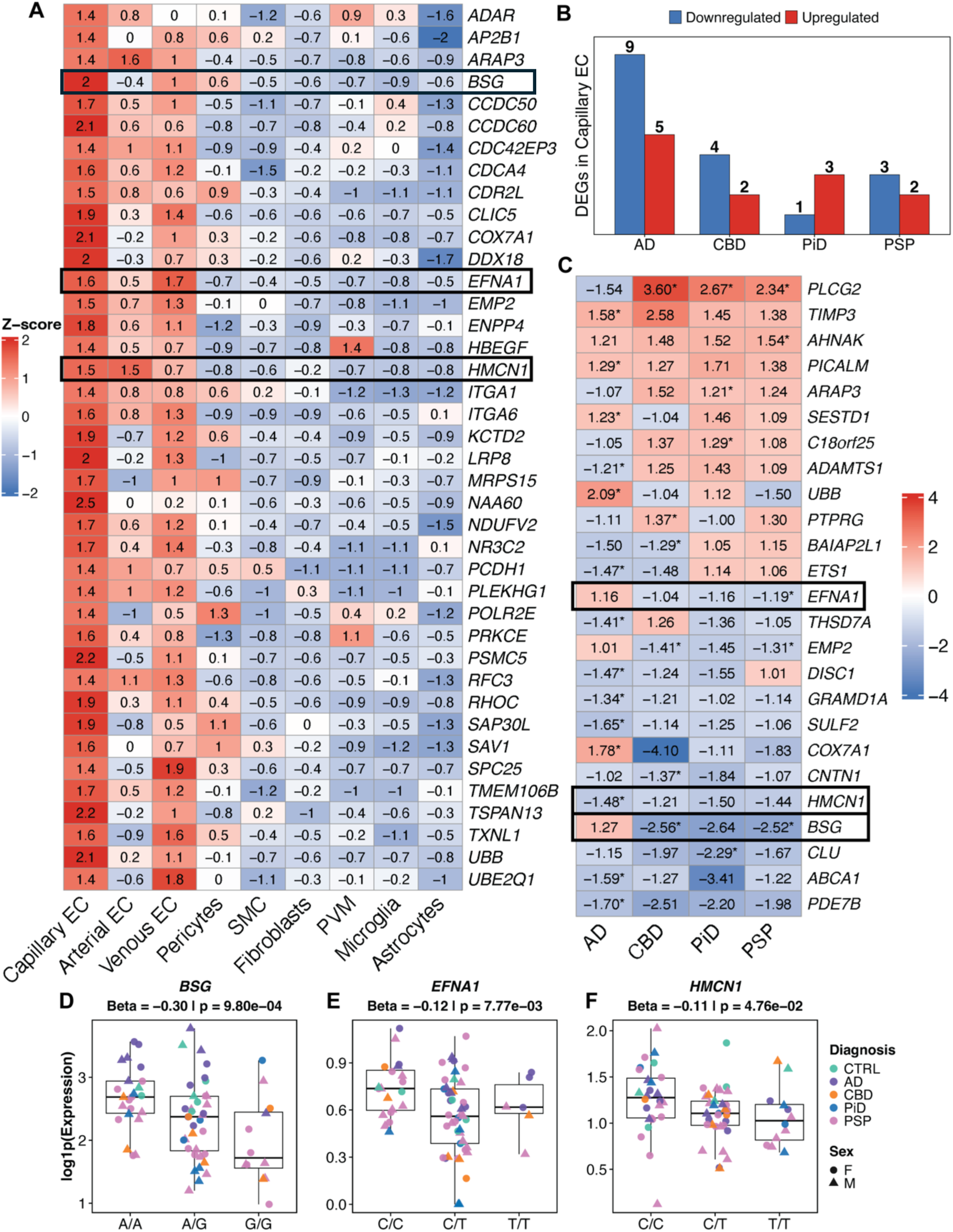
GWAS gene expression changes and polymorphic genetic influences within vascular-related cells. A) Heatmap displaying the top 40 AD GWAS genes (from 651 reported by Yang *et al.*) ranked by highest enrichment in capillary endothelial cells (Capillary ECs). Values represent z-scored average log1p expression across cells within each cell type. B) Bar plot summarizing the number of AD GWAS genes that are differentially expressed in capillary endothelial cells in each diagnosis vs control comparison. C) Heatmap showing scaled fold-change values for all 25 AD GWAS genes overlapping with capillary endothelial cell differentially expressed genes identified in at least one diagnosis vs. control comparison. Genes are ordered by mean scaled fold change across all four diagnoses. Asterisks indicate significant differential expression (MAST) in the corresponding diagnosis comparison. D-F) Boxplots showing donor-level log1p expression for the three genes from C) (BSG (D), EFNA1 (E) and HMCN1 (F)) with significant eQTL associations between genotype dosage and expression. eQTL analysis was performed using a linear model (Expression ∼ Dosage + Age + Sex + Diagnosis). Points represent individual donors, colored by diagnosis and shaped by sex.

Having noted that the AD risk gene *PLCG2* was upregulated in capillary endothelial cells of primary tauopathies, but not AD capillary endothelial cells, we next examined whether AD-risk genes are dysregulated across all diseases. As expected, the largest number of dysregulated AD-risk genes was observed in AD (14 genes). However, a small number of other genes were also significantly up- and down-regulated in CBD (6 genes), PiD (4 genes), and PSP (5 genes) (**Figure 5B**). Many of these gene changes were unique to only one disease (**Figure 5C**). In sum, these results highlight divergent transcription level changes in capillary endothelial cells within a subset of genes associated with AD risk.

To better understand how genetic polymorphisms could be influencing the transcriptional differences that were observed in this cell population, we performed eQTL analysis using genomic data from these donors. We identified three single-nucleotide polymorphisms (SNPs) that were significantly associated with expression level differences in capillary endothelial cells involving *BSG*, *EFNA1*, and *HMCN1*, which were all associated with reduced expression of these genes (**Figure D-F**). Altogether, these findings indicate that variations in these loci could be contributing to cell-type specific differences in gene expression that affect disease risk.

## Discussion

In this work, we systematically compared the relative contributions of cerebrovascular cell types across AD, CBD, PiD, and PSP. Our snRNA-seq analysis identified capillary endothelial cells and pericytes as the most transcriptionally perturbed vascular populations in the inferior temporal gyrus, with consistent upregulation of heat shock pathways across diseases and cell types, suggesting a shared cerebrovascular stress response. In contrast, AD exhibited a distinct transcriptional profile, characterized by broader gene expression changes and increased vascular remodeling, distinguishing it from primary tauopathies.

Transcriptomic data indicated that AD cases display increased capillary endothelial cell proportions, compared to primary tauopathies. We validated this observation using clear tissue immunolabeling on 1 mm-thick tissue sections from the same donors used for the transcriptomic analysis. While our sample set did not include enough tissues from control individuals to compare tauopathy to normal healthy aging, the number of vascular segments and total length were significantly reduced in tauopathies compared to AD. Qualitatively, the cerebrovasculature in primary tauopathies appeared less elaborate with fewer small diameter vessels, particularly in deeper areas of cortex and white matter. A prior study that included 11 PSP donors similarly reported a reduction in endothelial cell clusters^37^. Since white matter areas were excluded from our snRNA-seq dataset (via dissection), future studies evaluating white matter vasculature may reveal additional disease-relevant transcriptional changes not described here. Our findings showing increased cerebrovascular complexity in AD are in line with prior reports that used other methods to visualize vascular networks in human brain and have reported increased vascular density, length, and branching in AD^38–41^. However, it should be noted that others have seen opposite trends in these measures^42–44^. These discrepancies could be associated with differences in regional responses to disease or interindividual variability – factors not addressed by with our limited histology dataset. Importantly, Biron *et al.* reported increased microvessel density and suspected angiogenesis in the hippocampus and cortex of a mouse model of amyloid β pathology, which was validated in the cortex of *postmortem* human AD tissue^38^. In addition, Desai *et al.* showed that the hippocampus of human AD cases displayed increased vessel density that directly correlated with amyloid β plaque density, but not neurofibrillary tangles^39^. Together with our findings, the data suggest that the observed increase in vascular density in AD may be driven by amyloid β pathology, as we did not observe this in the primary tauopathy cases.

Aberrant angiogenesis can lead to immature vessel formation and result in faulty BBB integrity, which is well reported to occur in the early stages of AD^45,46^. Interestingly, increased cell proportions of pericytes and PVMs were found in primary tauopathies, compared to AD. Evidence of pericyte loss and dysfunction has been reported in AD^3,47^; however, the potential implications of increased pericyte and PVM cell populations in primary tauopathies remain widely unknown. Under pathological stress conditions, pericytes can modulate neuroimmune responses within the CNS and facilitate the entry of peripheral immune cells across the BBB^48^. Related to AD, amyloid β oligomers have been shown to target and contribute to pericyte cell dysfunction, which could lead to changes in cerebral blood flow^49,50^. While our data showed comparable pericyte cell populations between AD and control cases, we found that primary tauopathies with relatively absent amyloid β pathology display increased pericyte cell populations. These findings highlight the potential influence of amyloid β on cerebrovascular pericytes and could explain the drastic changes in cerebral blood flow that take place in these conditions.

PVMs have also been shown to regulate peripheral immune cell infiltration into the central nervous system and have been reported to accumulate ROS that can damage the neurovascular unit^51,52^. Peripheral immune cell infiltration in AD and related tauopathies remains a controversial research topic. However, convincing reports have shown the infiltration of T-cells across the BBB into the brain parenchyma of both AD brains and brains of tauopathies without amyloid beta aggregation^53,54^. Importantly, not amyloid β, but rather tau pathology was found to significantly correlate with T-cell infiltration in cases in which both pathologies were present^55^. Our data here indicate that pericyte and PVM cell populations increase in the presence of tau pathology, but that this may be halted by amyloid β pathology in AD.

In addition to cerebrovascular remodeling, the upregulation of heat shock proteins across disease states suggests a common stressor. The heat shock protein family is comprised of molecular chaperones that fold peptides, degrade misfolded proteins, and inhibit aggregation. In our data, *HSP90AA1* was upregulated in all diseases, and this was directly translated to the protein level. *HSP90AA1* encodes the protein HSP90α and is induced in cells under stress, unlike the constitutively active *HSP90AB1*^56^. While HSP90α can interact with an estimate 10% of all human proteins, it notably interacts with tau and can impact its degradation^57,58^. Prior studies have shown that endothelial cells are capable of taking up misfolded tau, and this may be one explanation for the universal upregulation of this protein across diseases^59,60^.

Other gene expression signatures that were also unique to AD capillary endothelial cells involve enhanced ribosome biogenesis, which is important for cell migration and proliferation, and increased expression of immune signaling molecules. Interestingly, the upregulation of *HLA-B* in AD endothelial cells could indicate that these cells are acting as antigen presenting cells and interacting with CD8^+^ T-cell populations, which have been implicated in AD pathogenesis. By contrast, *HLA-A,* which performs similar functions, appeared to be reduced in tauopathy.

However, evaluation by histology showed that protein expression may not fully recapitulate changes seen at the gene expression level. Additional studies in a larger histological cohort, which includes co-labeling for T cell populations, would be required to confirm these relationships. Similarly, increases in cysteine-rich metal ion binding proteins – metallothioneins – suggests a response to oxidative stress not observed in tauopathies. Last, increased *CLDN5* expression in AD was unexpected based on prior reports of BBB disruption in AD but indicates that disruption may be occurring via mechanisms other than depletion of tight junction proteins. This finding is consistent with an AD proteomics study that similarly observed increased CLDN5^61^.

Lastly, genes associated with AD-risk were notably enriched in capillary endothelial cells and overlapped to the greatest extent with AD DEGs, confirming a role for these cell types in influencing disease progression. However, AD-risk genes were also significantly altered in tauopathies and may reveal divergent roles for endothelial cells in neurodegeneration. By evaluating cell-type eQTLs in this dataset, we identified SNPs that impact expression of three AD-risk associated genes. *BSG and EFNA11* SNPs were associated with reduced expression of these genes in tauopathy. *BSG* encodes the protein Basigin (CD147), which is involved in VEGF signaling and *EFNA1* encodes an EphA2 receptor ligand implicated in endothelial cell migration and tumor growth^62–65^. The downregulation of both genes in tauopathy indicates a suppression of pro-angiogenesis pathways that could help explain the relative reduction in capillary endothelial cells observed. A SNP in *HMCN1,* which encodes the extracellular matrix protein Hemicentin, was associated with reduced expression in AD. In our prior work, *HMCN1* was also a marker gene of capillary endothelial cells in primary visual cortex, a region that develops limited AD pathology, indicating that *HMCN1* may have different roles in other brain areas^12^. Further exploration would be needed to understand the influence of this protein on disease.

Our findings highlight fundamental differences in cerebrovascular alterations between AD and primary tauopathies. While neuroimaging studies have described reduced cerebral blood flow as an early feature common to dementias^7,43^, our work indicates that the underlying cause of cerebral blood flow reductions are unlikely to be shared mechanisms across diseases. Rather, these results indicate that diverse transcriptional programs impact vascular structure and function. By contrast, heat shock protein upregulation appears to be a shared biological process among the neurodegenerative conditions studied here. Our results warrant future studies that evaluate heat shock responses in other neurodegenerative diseases such as Huntington’s and Parkinson’s disease to determine whether these are general responses of vascular cells to aggregated protein stressors, or a specific feature of tauopathies. Overall, by selectively enriching vascular cells and using methods to evaluate brain vasculature, this work reveals significant changes in transcriptional and translational signatures, vascular remodeling, and genetic polymorphic influences within these cell types in AD and primary tauopathies. Understanding how these vascular changes contribute to disease progression and pathogenesis may enable the development of more precise, mechanism-informed therapeutic strategies targeting cerebrovascular dysfunction in neurodegeneration.

## Data Availability

The snRNA-seq dataset generated in this study will be made available at Gene Expression Omnibus (GEO: GSE327289) upon acceptance of the paper. Raw sequencing data (FASTQ files) for a subset of samples have been previously deposited in the Sequence Read Archive (SRA) under PRJNA916657. Due to ongoing analyses and related unpublished studies, raw FASTQ files for the remaining samples are not publicly available at this time.

## Code Availability

All code generated in this study is available at https://github.com/mindds/Vascular-AD-Tauopathy. All software used in this study is publicly available.

## Author Contributions

RB, SD, LDE, SD, AG, AD, TK, MW, RVT, and YG conceived of and designed the study. LDE, SD, AG, SC, TP, AA, JT, KY, NR, KZ, AM, EA, TK and YG performed the experiments. LDE, SD, AG, SY, PPS, XW, GL, and EA analyzed the data. BTH, TJZ, ASP, DO, FL, LG, JR, RT, MW, YG provided critical feedback and guidance for experimental and analysis design. LDE, SD, AG, SD, and RB wrote the manuscript. All authors read and approved of the final manuscript.

## Ethics Declarations

MW, YG, SC, KY, NR, KZ, SY, PPS, XW, GL, AM, EA, AD, JR are employees of AbbVie. AA, TP, JT, TK, and RVT were AbbVie employees at the time of the study. The design, study conduct, and financial support for this research were provided by AbbVie.

## Acknowledgements

ASP, BTH and SD were supported by the NIH/NIA P30AG062421 (Massachusetts Alzheimer’s Disease Research Center). TJZ supported by Coins for Alzheimer’s Research Trust. RB supported by the NIH/NIA R01AG071567.

---

**Supplemental Figure 1.**
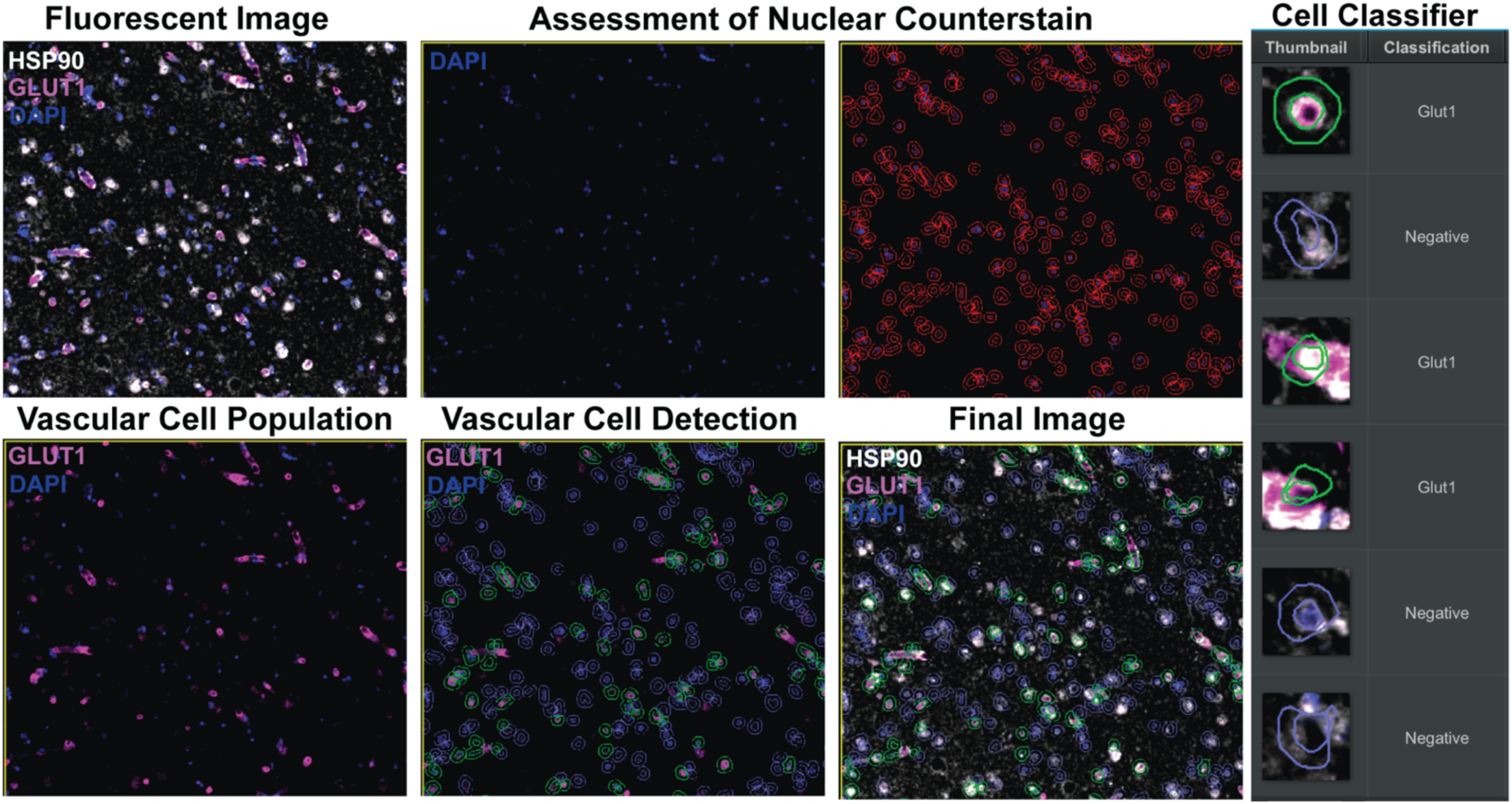
Quantification of protein markers within individual GLUT1^+^ cells. Human ITG (BA20) tissue sections were stained with fluorescent markers of interest. Using a VS120 microscope slide scanner at 20x magnification, sections were imaged in the 405, 488, 591, and 647 nm fluorescent channels. Output scans were imported into Qpath pathology software (Version 0.6.0) in which DAPI-positive vascular cells were detected by applying the program’s object classifier based on staining intensity (nuclear counterstain assessment). The classifier data was exported to record the mean intensity of each individual marker of interest within each individual GLUT1^+^ cell.

**Supplemental Figure 2.**
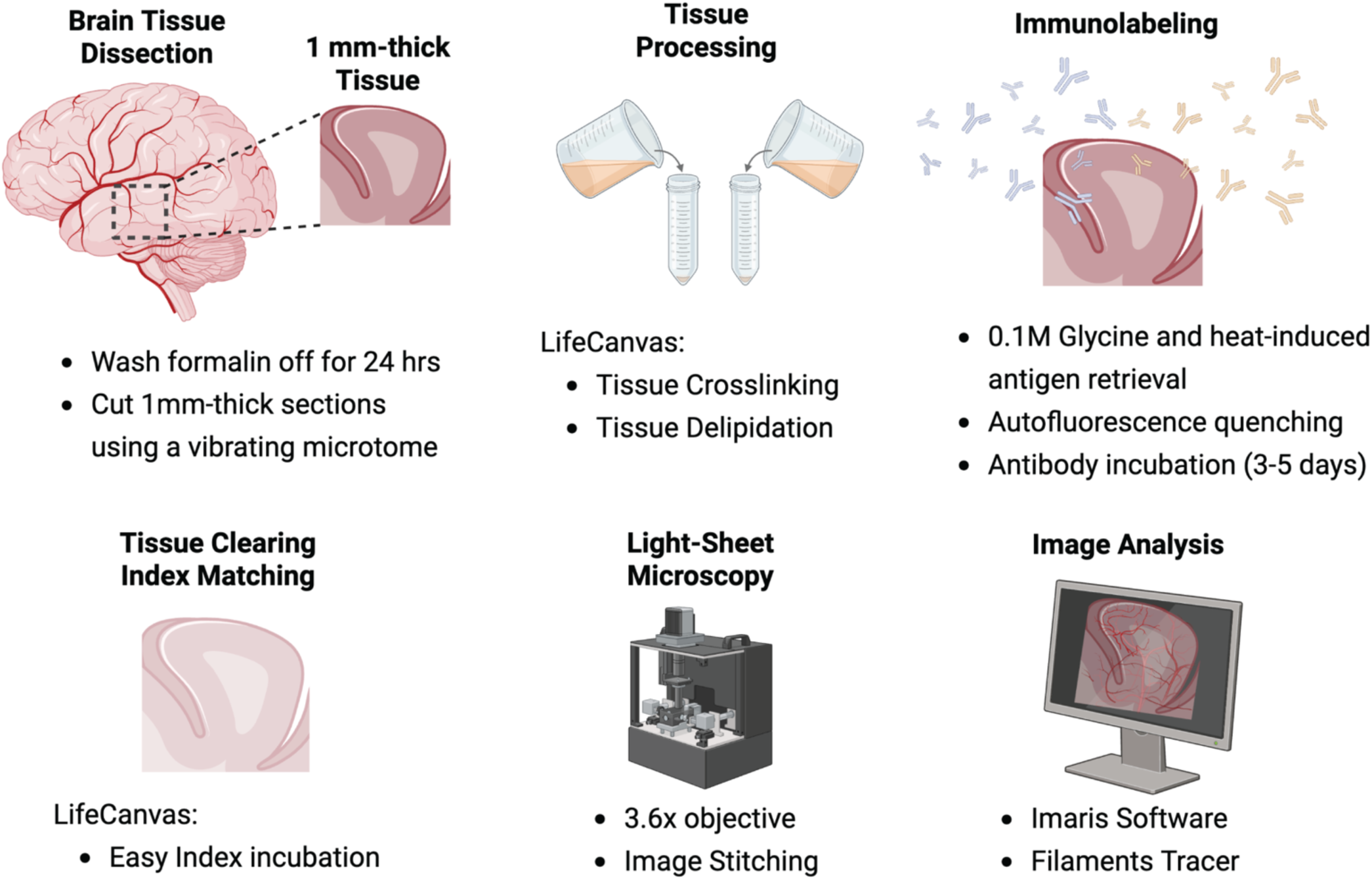
Application of tissue clearing techniques to formalin-fixed ITG samples. Summarized graphical workflow of tissue processing starting from formalin-fixed tissue dissection until light-sheet microscopy and image analysis. Created in Biorender.com

**Supplemental Figure 3.**
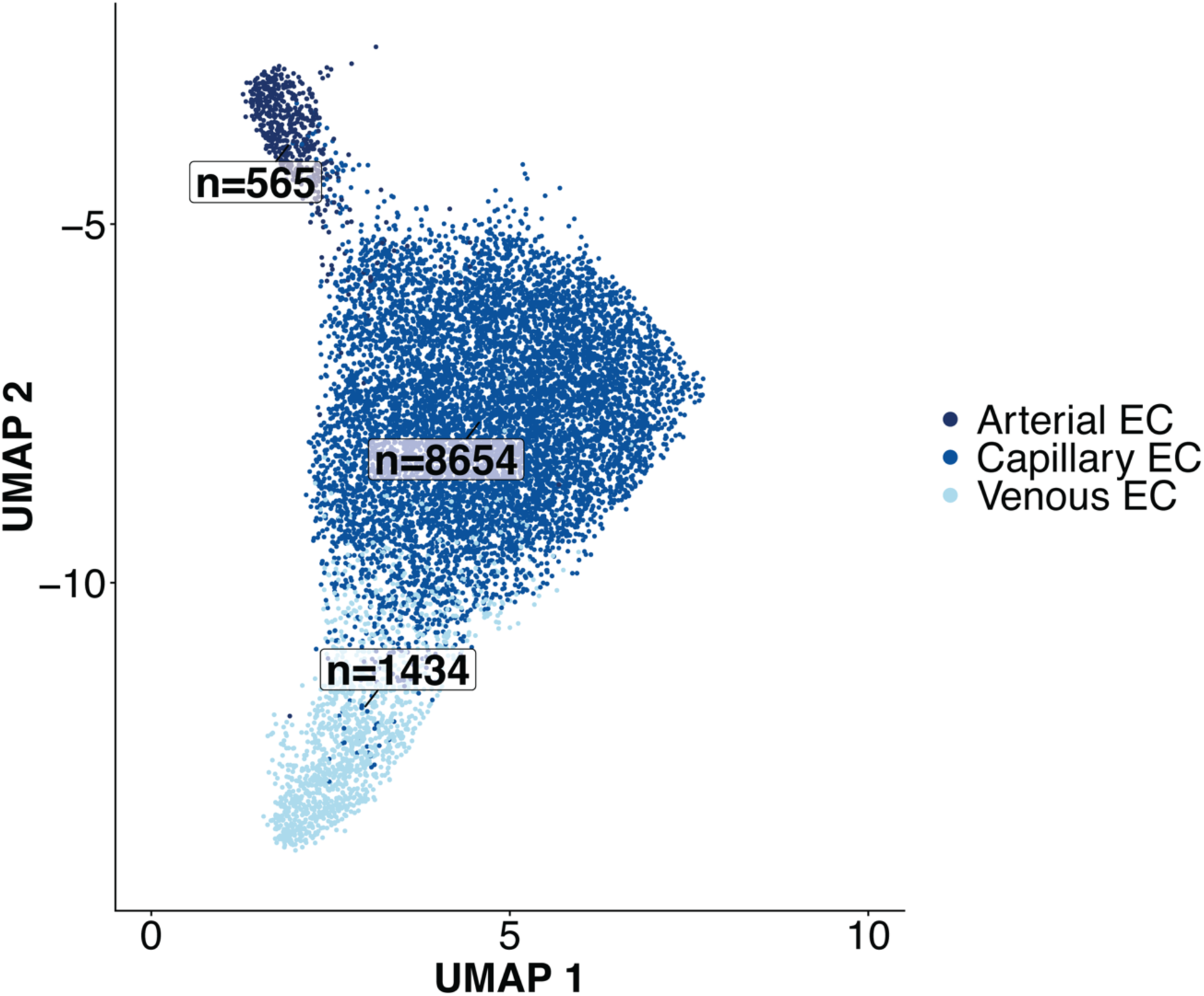
UMAP visualization of the identified endothelial cell clusters, each containing their corresponding number of identified nuclei.

**Supplemental Figure 4.**
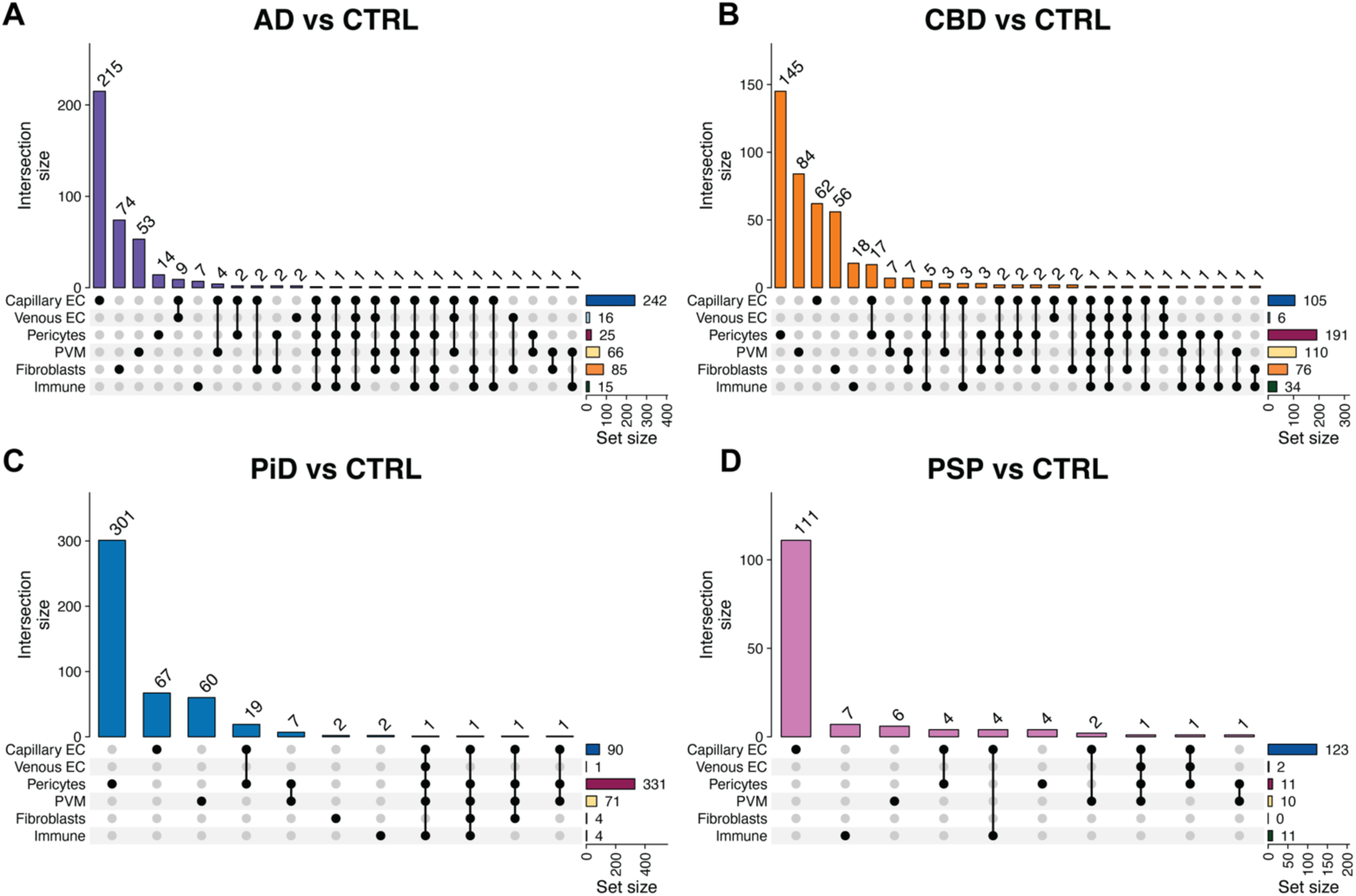
A) – D) Upset plots displaying the number of DEGs for each identified cell type and whether DEGs are shared between such comparisons in AD (A), CBD (B), PiD (C), and PSP (D).

**Supplemental Figure 5.**
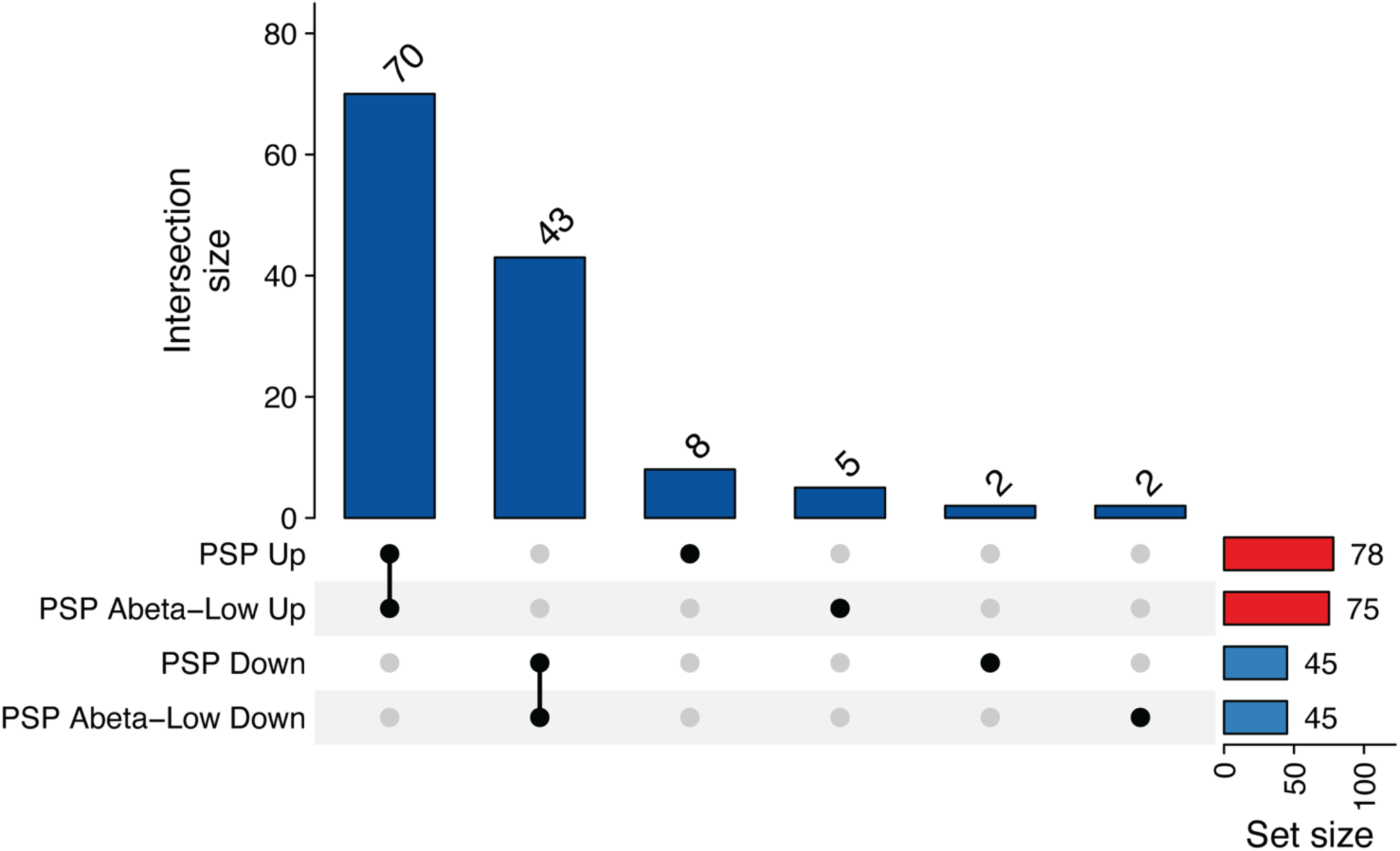
Sensitivity analysis confirming that PSP vs. Control DEGs in Capillary Endothelial Cells are independent of amyloid beta burden. UpSet plot showing the overlap between DEGs from PSP vs. Control and PSP with low amyloid beta burden vs. Control comparisons, separated by direction of effect (up- and downregulated). Low amyloid beta burden was defined as less than 1% amyloid beta percent area.

